# Canonical Wnt Signaling Suppresses Brain Endothelial Cell Transcytosis to Maintain Blood-Brain Barrier Integrity

**DOI:** 10.1101/2025.03.10.642437

**Authors:** Xavier du Maine, Chenghua Gu

## Abstract

Canonical Wnt signaling is essential for blood-brain barrier (BBB) development and maintenance. However, the subcellular mechanisms underlying this critical regulation have remained elusive. In this study, we use a physiological paradigm examining an early phase of acutely attenuated canonical Wnt signaling in adult brain endothelial cells (ECs) to investigate how the pathway regulates BBB integrity. Following canonical Wnt signaling attenuation via EC-specific knockout of β-catenin, we find that there is increased transcytosis in brain ECs, including a striking diversity of morphologically distinct vesicles, indicating multiple pathways are involved. In addition, we find that although the molecular composition of tight junctions (TJs) is altered following canonical Wnt signaling attenuation, such that Claudin-5 and ZO-1 expression is downregulated, TJs remain impermeable to molecules as small as 1.9 kDa. These findings reveal previously underappreciated role of Wnt signaling in regulating brain EC transcytosis and help illuminate subcellular mechanisms of BBB maintenance in adulthood, which is crucial for improving delivery of therapeutics to the brain.

## Introduction

The blood-brain barrier (BBB) maintains a homeostatic environment for optimal neuronal function by tightly regulating the movement of ions, molecules, and cells between the blood and brain parenchyma. The primary site of the BBB occurs at brain endothelial cells (ECs) that form tightly sealed blood vessels. At the subcellular level, these ECs have specialized properties that determine BBB structural integrity: highly impermeable tight junctions (TJs), suppression of transcellular vesicular trafficking (transcytosis), and the absence of fenestrations^1,2^.

An essential regulator of the BBB is canonical Wnt signaling, which orchestrates a transcriptional program in brain ECs that is dependent on the transcriptional co-activator β-catenin^3^. The genes upregulated by this pathway include some that are known to regulate BBB integrity, like the TJ protein Claudin-5 (CLDN5)^4^, as well as MFSD2A^5,6^, which suppresses transcytosis by inhibiting caveolae formation at the BBB^7,8^. During embryonic development, canonical Wnt signaling facilitates brain angiogenesis and barriergenesis^9-11^. In addition, early postnatal brain angiogenesis, as well as maintenance of BBB integrity during early postnatal development and adulthood, all require active canonical Wnt signaling^12-14^. Accordingly, mouse models with loss of function mutations in the genes encoding brain-expressed Wnt ligands^13,15^ or brain EC-expressed Wnt receptors^12^, co-receptors^13^, co-activators^16,17^, or β-catenin^13^, display increased BBB permeability to small molecular weight exogenous tracers and/or large endogenous serum proteins, suggesting an increase in paracellular and/or transcellular permeability.

Despite this large body of research, the subcellular mechanisms that govern canonical Wnt signaling regulation of BBB integrity remain elusive. First, evaluation of TJs in various Wnt pathway mutants has been largely limited to TJ protein expression levels, establishing correlative but not causal relationships between TJ protein levels and BBB permeability^9,12,13,15,18^. Critically, there have been no functional studies directly evaluating TJ permeability using tracer-coupled electron microscopy (EM), which is required for distinguishing paracellular from transcellular permeability at the subcellular level. Second, the contribution of transcytosis has remained underexplored *in vivo* by EM (or other analysis with subcellular resolution) despite the identification of MFSD2A as a Wnt-regulated gene by microarray, RNA-seq, and immunohistochemical analyses^5,6,18^.

These subcellular mechanisms have been difficult to ascertain due to the pathway’s confounding roles in different physiological and pathological processes. First, canonical Wnt pathway mutants have severe CNS angiogenesis defects^5,10,11,13,16,17,19-21^. Therefore, when using these mutants, especially at embryonic and early postnatal stages, it is not possible to rule out if changes in subcellular properties are due to broader effects of a developmentally immature vasculature^22^. Second, within 1-2 weeks following acute EC-specific ablation of β-catenin in adult mice, there are a host of neurological pathologies, including abundant neuronal cell death, petechial hemorrhages, and lethal seizures^23^. Therefore, at this time point, it is not possible to rule out if changes in subcellular properties are secondary to these pathologies.

In this report - using a combination of EC-specific mouse genetics, BBB leakage assays, and tracer-coupled quantitative EM - we circumvent these confounds by using a physiological paradigm to examine an early phase (3 days after tamoxifen injections) of acutely attenuated canonical Wnt signaling in adult brain ECs. Specifically, we acutely knock out β-catenin in ECs of adult mice. In doing so, we observe widespread BBB leakage and an increase in transcytosis, including a striking diversity of vesicular morphologies, while TJ impermeability is not overtly compromised, demonstrating that suppression of diverse pathways of transcytosis is a critical function of canonical Wnt signaling in brain ECs.

## Results and Discussion

### A physiological paradigm of acutely attenuated canonical Wnt signaling in adult brain ECs

In order to acutely attenuate canonical Wnt signaling in brain ECs, we knocked out the gene encoding β-catenin (*Ctnnb1*) by crossing *Ctnnb1* floxed mice^24^ (*Ctnnb1*^*fl/fl*^) to EC-specific tamoxifen-inducible *Cdh5-Cre*^*ERT2*^ mice^25^. To circumvent the confounding angiogenesis defects at earlier developmental time points, we performed five consecutive days of tamoxifen injections in adult mice (8-10 weeks), when brain ECs are largely quiescent^26^ (Figure 1a). Since complex neurological pathologies were observed 1-2 weeks following tamoxifen treatment^23^, we assayed mice three days following the last tamoxifen injection, well before the onset of seizure-associated pathology (Supplementary Figure 1a). At this time point, mutants (*Cdh5-Cre*^*ERT2*^*;Ctnnb1*^*fl/fl*^) did not exhibit cell death or decreases in vessel density (Figure 1b-d). To assess canonical Wnt signaling in brain ECs, we immunostained for the transcription factor LEF1, which is an effector and transcriptional target of the canonical Wnt pathway in ECs^27^. We observed a ∼40% reduction in LEF1 intensity in brain EC nuclei in mutants relative to controls (*Ctnnb1*^*fl/fl*^) across multiple brain regions (Figure 1e-f, Supplementary Figure 1b-c). Together, these results demonstrate that we were able to acutely attenuate canonical Wnt signaling in adult brain ECs without inducing pathology.

**Figure 1.**
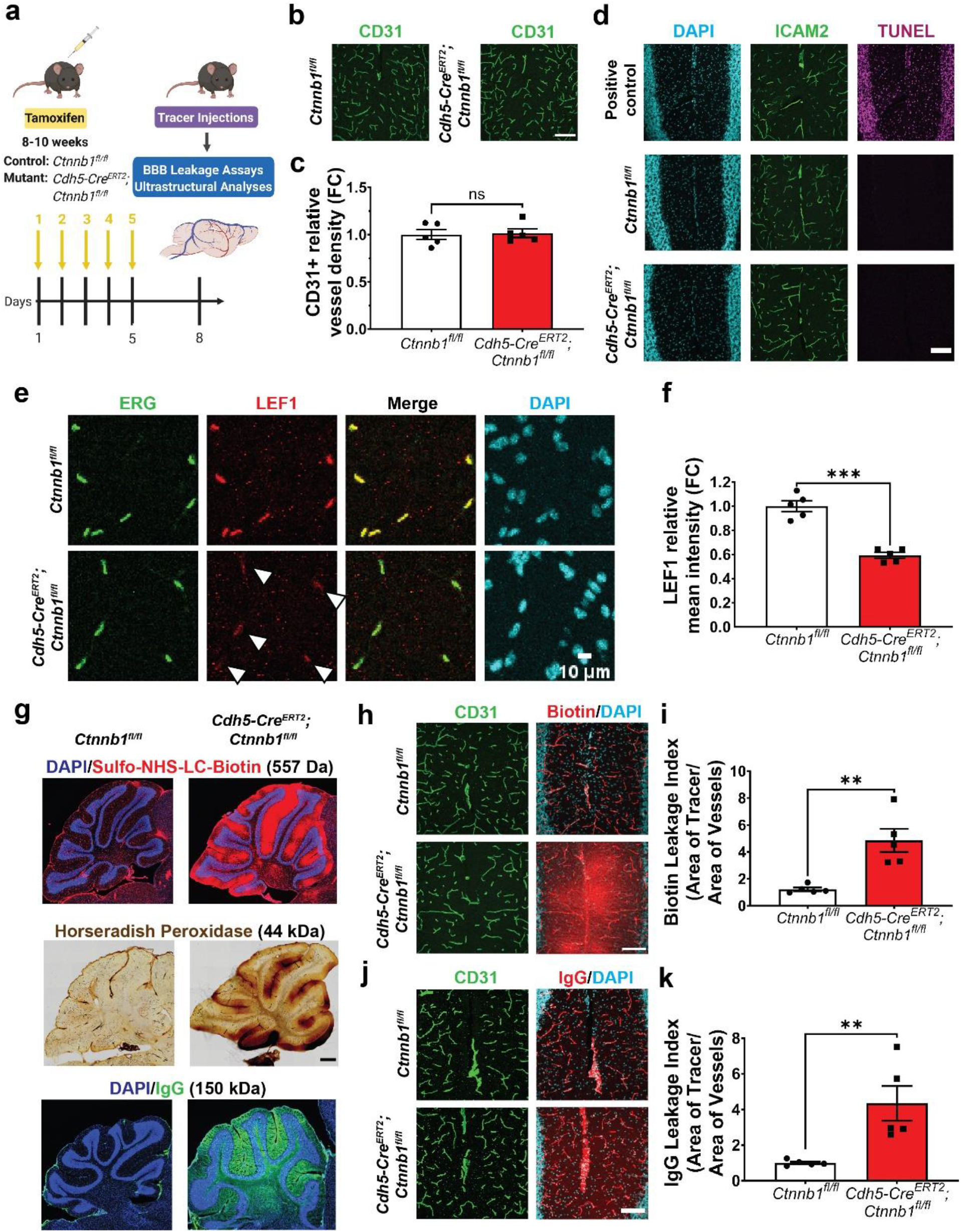
Early-phase acute attenuation of canonical Wnt signaling in the adult brain vasculature results in BBB leakage. **(a)** Schematic overview of experimental paradigm: all animals were injected with tamoxifen for 5 consecutive days and assays were performed 3 days after the last injection. Created with BioRender.com **(b)** Immunofluorescence images of brain ECs (CD31) in controls (*Ctnnb1*^*fl/fl*^) and mutants (*Cdh5-Cre*^*ERT2*^*;Ctnnb1*^*fl/fl*^). (Scale bar: 100 µm) **(c)** Quantification of vessel density (data are mean ± SEM, n=5 animals per genotype). ns = not significant, unpaired t test. **(d)** TUNEL staining (magenta) in the cerebellum of control (middle) and mutant (bottom) mice, as well as DNase-treated positive control (top), with vessel (ICAM2) and nuclear (DAPI) staining. (Scale bar: 100 µm) **(e)** Immunofluorescence images of EC nuclei (ERG), canonical Wnt signaling marker LEF1, and all nuclei (DAPI) in the cerebellar molecular layer of control and mutant mice. White arrowheads denote reduced LEF1 staining in mutant EC nuclei. (Scale bar: 10 µm) **(f)** Quantification of the LEF1 mean fluorescence intensity across ERG^+^ EC nuclei (data are mean ± SEM, n=5 animals per genotype). ***p<0.001, unpaired t test with Welch’s correction. **(g)** Immunofluorescence images of sulfo-NHS-LC-biotin (top) and IgG (bottom) BBB leakage in the cerebellum with nuclear (DAPI) counterstain. Brightfield image of horseradish peroxidase (middle) leakage in the cerebellum. Confocal images of sulfo-NHS-LC-biotin **(h)** and IgG **(j)** leakage in the cerebellar molecular layer with vessel (CD31) counterstain. (Scale bar: 100 µm) Quantification of the biotin **(i)** and IgG **(k)** leakage indices: area of tracer/area of CD31^+^ vessels (data are mean ± SEM, n=5 animals per genotype). **p<0.01, Mann-Whitney test.

To assess BBB integrity during this early acute phase of attenuated canonical Wnt signaling, we injected tracers into the blood circulation then determined if they leaked out of the brain vasculature. For our study, we assayed leakage of the small molecular weight tracer sulfo-NHS-LC-biotin (557 Da), the larger tracer horseradish peroxidase (HRP, 44 kDa), and the endogenous serum protein IgG (150 kDa). Biotin leakage was widespread, with the most severe leakage occurring in the cerebellum, hippocampus, and olfactory bulbs (Figure 1g-i and Supplementary Figure 1d). HRP leakage, though similarly distributed, was not as severe. Lastly, IgG leakage was confined to the cerebellum and olfactory bulbs (Figure 1g,j-k and Supplementary Figure 1d). Overall, the strongest leakage phenotypes occurred in the cerebellum and olfactory bulbs (Supplementary Figure 1d).

The occurrence of interregional heterogeneity in BBB leakage - despite a similar decrease in EC LEF1 levels across these regions (Supplementary Figure 1b-c) - suggests an increased sensitivity to attenuation of canonical Wnt signaling in the cerebellum and other similarly affected regions. These results raise multiple possibilities. First, expression levels of transcription factors downstream of β-catenin and LEF1 (e.g. SOX17, ZIC3, FOXQ1) may differ between ECs in different brain regions. Second, ECs in regions with less severe leakage may possess β-catenin-independent parallel pathways for maintaining BBB integrity in adulthood. Interestingly, the cerebellar BBB leakage in our study mirrors an earlier study of acute EC-specific loss of the upstream Wnt receptor FZD4 in adult mice, which resulted in BBB breakdown restricted to the cerebellum^12^. Due to the consistent and anatomically stereotyped nature of BBB leakage across multiple tracers in the cerebellum, and in particular the cerebellar molecular layer, we focused all subsequent analyses on this subregion.

### Early-phase acute loss of canonical Wnt signaling in the adult brain vasculature alters the molecular composition of TJs without overtly compromising impermeability

Having established an increased BBB permeability to exogenous tracers and endogenous serum protein, we next sought to identify the subcellular mechanisms underlying BBB leakage. We first evaluated the molecular composition of TJs in mutant mice, staining for CLDN5, the predominant component of brain EC TJs and a known transcriptional target of the pathway, as well as its cytoplasmic partner ZO-1. In mutant ECs, we observed a ∼40% reduction in CLDN5 levels (Figure 2a-b), and a very small but significant ∼10% reduction in ZO-1 levels (Figure 2c-d).

**Figure 2.**
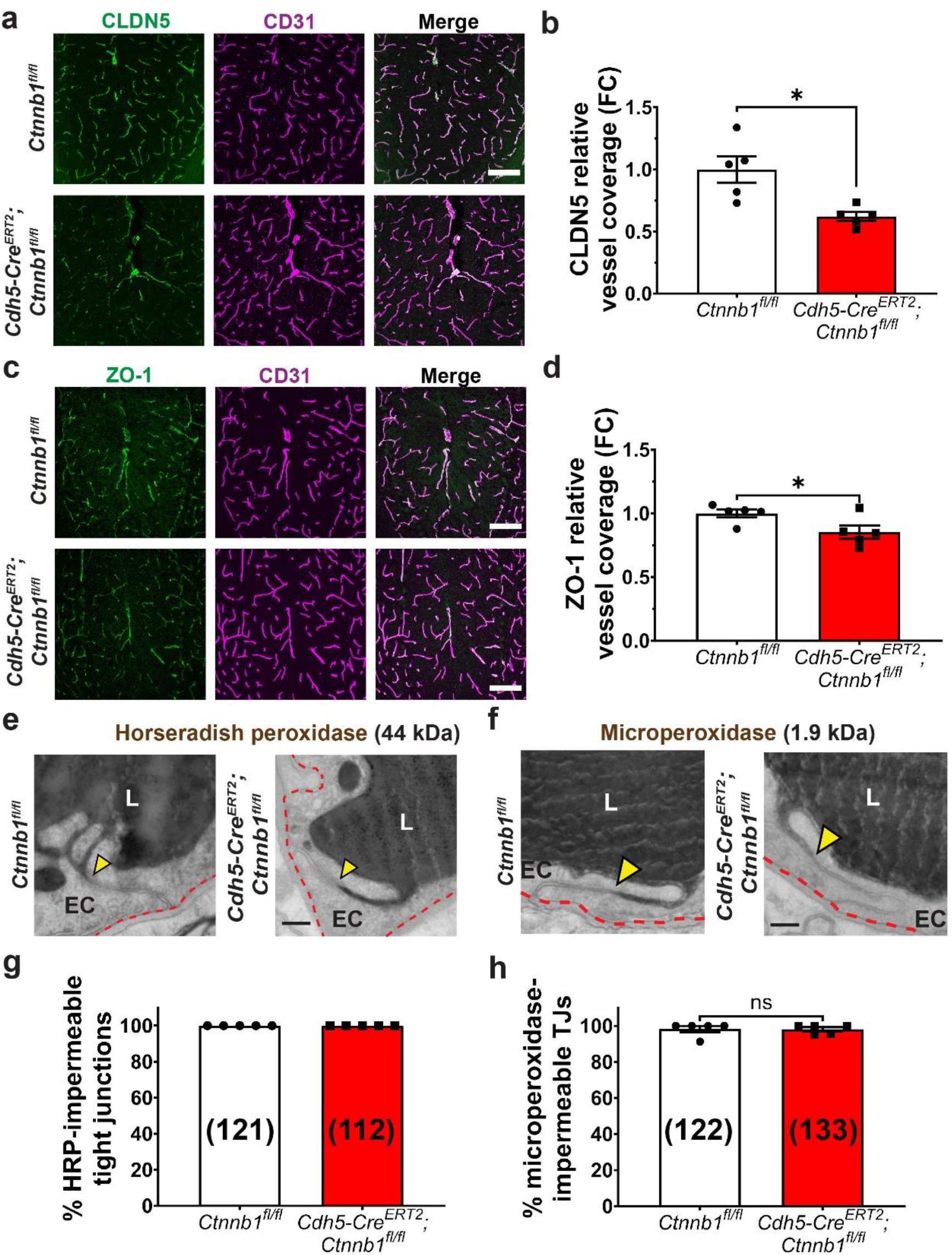
Early-phase acute attenuation of canonical Wnt signaling in adult brain ECs alters the molecular composition of TJs without affecting TJ permeability to tracers as small as 1.9 kDa. **(a)** CLDN5 immunofluorescence staining in the cerebellar vasculature with CD31 (EC) co-stain. Scale bar: 100 µm. **(b)** Quantification of CLDN5 relative levels normalized to controls (data are mean ± SEM, n=5 animals per genotype, *p<0.05, unpaired t test with Welch’s correction). **(c)** ZO-1 immunofluorescence staining in the cerebellar vasculature with CD31 (EC) co-stain. Scale bar: 100 µm. **(d)** Quantification of ZO-1 relative levels normalized to controls (data are mean ± SEM, n=5 animals per genotype, *p<0.05, unpaired t test). Electron micrographs showing cross sections of cerebellar ECs demonstrate that both HRP **(e)** and microperoxidase **(f)** are halted at TJs (yellow arrowheads) in both control (*Ctnnb1*^*fl/fl*^) and mutant (*Cdh5-Cre*^*ERT2*^*;Ctnnb1*^*fl/fl*^) mice. Scale bar: 200 nm. **(g)** Quantification of data in (e) showing that 100% of TJs observed were impermeable to HRP (n=5 animals per genotype, total number of TJs observed per genotype in parentheses). **(h)** Quantification of data in (f), showing that the vast majority of TJs observed in control (*Ctnnb1*^*fl/fl*^) and mutant (*Cdh5-Cre*^*ERT2*^*;Ctnnb1*^*fl/fl*^) mice were impermeable to microperoxidase (data are mean ± SEM, n=5 animals per genotype, total number of TJs per genotype in parentheses, ns = not significant, unpaired t test). L = lumen, EC = endothelial cell. Red dashed lines outline abluminal membranes of ECs.

To examine the functionality of TJs at this early phase, we evaluated TJ permeability using electron microscopy (EM) in HRP-injected mice. HRP has an electron-dense reaction product in the presence of its substrate DAB that appears as black in electron micrographs, enabling its subcellular visualization in ECs. As a positive control for permeable TJs, we used choroid plexus ECs, which have very low Wnt signaling and constitute continuous fenestrated vessels^18^. In the choroid plexus, we observed leakage of HRP and the smaller tracer microperoxidase (1.9 kDa) through endothelial TJs (Supplementary Figure 2). In contrast, in all control and mutant cerebellar ECs, we observed that HRP was sharply halted at TJs, indicating that TJs were impermeable to HRP (Figure 2a-b). To detect more subtle changes in TJ permeability, we employed microperoxidase. Like HRP, microperoxidase was also sharply halted at TJs, with nearly 100% impermeable to this tracer (Figure 2c-d).

Given that in previous studies^9,12,13,15^ CLDN5 expression has been anticorrelated with expression of PLVAP – a structural component of fenestral diaphragms^28,29^ – we next stained for PLVAP. In the cerebellum, we found that PLVAP expression was overall very sparse and spatially restricted to the central axis of the molecular layer (Supplementary Figure 3a), in contrast with the widespread BBB leakage observed in mutants (Figure 1, Supplementary Figure 1d). The small magnitude and short timescale of this PLVAP expression is unlikely to correlate with *de novo* fenestration formation. In accordance with this postulation, across the 509 vessels we assessed in mutant animals in this study, we observed no fenestrations (Figure 2e-h, Figure 3a-c,f-h).

**Figure 3.**
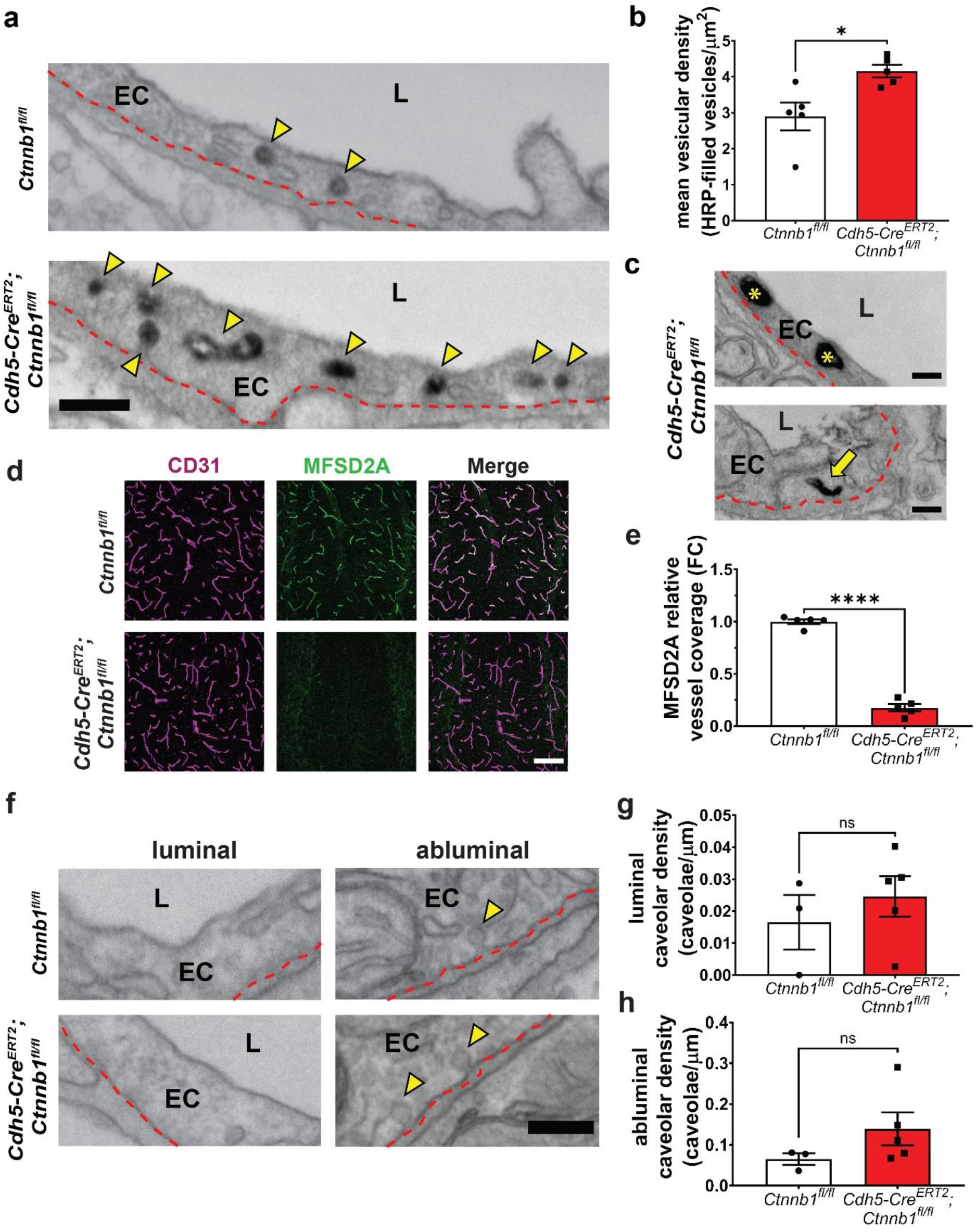
Suppression of transcytosis by canonical Wnt signaling is required for blood-brain barrier integrity. **(a)** Representative EM high magnification images of cerebellar molecular layer ECs following HRP (black) injection and low-pressure gravity perfusion. Yellow arrowheads show HRP-filled vesicles, red dashed lines denote the abluminal membrane of ECs; EC=endothelial cell, L=lumen (Scale bar: 200 nm) **(b)** Quantification of mean vesicular density (number of HRP-filled vesicles per µm^2^ of EC cytoplasm): data are mean±SEM, n=5 animals per genotype (125 control vessels and 125 mutant vessels), *p<0.05, nested t-test. **(c)** Representative high magnification images of HRP-filled large spherical and tubular vesicles in mutant mice. (Scale bar: 200 nm) **(d)** Immunofluorescence images of MFSD2A in the cerebellar molecular layer with vessel (CD31) counterstain. (Scale bar: 100 µm) **(e)** Quantification of relative MFSD2A levels as fold change relative to control (data are mean±SEM, n=5 animals per genotype). ****p<0.0001, unpaired t test. **(f)** Representative EM high magnification images of luminal (left) and abluminal (right) caveolae vesicles in cerebellar molecular layer ECs. Note the rare occurrence of luminal caveolae. Yellow arrowheads show caveolae, red dashed lines denote the plasma membrane of ECs; EC=endothelial cell, L=lumen (Scale bar: 200 nm) **(g**,**h)** Quantification of luminal (g) and abluminal (h) caveolar density: data are mean±SEM, n=3 or 5 animals/genotype (96 control vessels and 139 mutant vessels), ns=not significant, nested t-test.

Together, these data demonstrate that although acute attenuation of canonical Wnt signaling in adult brain ECs alters the molecular composition of TJs, it does not alter TJ permeability to macromolecules as small as 1.9 kDa. The surprising contrast between these molecular and functional phenotypes underscores the need to evaluate TJ function using tracer-coupled EM rather than using TJ protein levels or overall TJ ultrastructure as a proxy for TJ integrity, which can lead to erroneous findings^22,23,30^. Notably, these TJ phenotypes are in accordance with the previously reported size-selective BBB leakage of *Cldn5*^*-/-*^ mice, which display leakage of smaller molecular weight tracers like sulfo-NHS-biotin (443 Da) but not larger tracers like microperoxidase^4^. While we found that mutant TJs are impermeable to HRP and microperoxidase, it is possible that they are permeable to smaller molecules like sulfo-NHS-LC-biotin, for which we observed severe BBB leakage (Figure 1g-i). The development and optimization of robust small molecule EM-compatible tracers to evaluate TJ permeability will enable these mechanistic investigations.

### Suppression of diverse types of vesicular-mediated transcytosis by canonical Wnt signaling is required for blood-brain barrier integrity

We next evaluated transcytosis using EM in HRP-injected mice with our paradigm of acute attenuation of canonical Wnt signaling in adult brain ECs. In mutant mice, there was a ∼40% increase in the density of HRP-filled vesicles in cerebellar ECs (Figure 3a-b). Notably, we observed a diversity of vesicular morphologies, including large spherical vesicles as well as tubular vesicles, indicating a possible upregulation of multiple vesicular pathways (Figure 3c). Furthermore, we observed HRP accumulation in the basement membrane (Supplementary Figure 3b-c), providing ultrastructural evidence of HRP leakage across the BBB. Given our earlier findings that mutant EC TJs were impermeable to HRP and that there were no fenestrations, these data collectively demonstrate that HRP leakage in mutant mice is due to increased transcytosis and that canonical Wnt signaling suppresses transcytosis to maintain BBB integrity. Moreover, given that mutant TJs are impermeable to HRP (44 kDa), we postulate that leakage of the much larger serum protein IgG (150 kDa) is likely also solely due to increased transcytosis. β-catenin’s role as an adherens junction protein is highly unlikely to contribute to the observed BBB leakage due to the fact that even molecules as small as microperoxidase (1.9 kDa) are unable to breach mutant junctions, indicating that overall vessel structural stability is unperturbed.

To evaluate the molecular mechanism underlying the increased transcytosis phenotype, we examined levels of MFSD2A in control and mutant mice by immunohistochemistry (Figure 3d). In mutant mice, there was a robust ∼90% decrease in MFSD2A protein levels (Figure 3e). Given this phenotype, we hypothesized that increased transcytosis in mutant mice may be mediated in part by MFSD2A loss and a resulting increase in caveolae - mediated transcytosis. To test this hypothesis, we quantified luminal and abluminal caveolae vesicles in mutant mice (Figure 3f). Strikingly, although we found that there was a trend toward an increase in caveolae, there was no significant difference in luminal or abluminal caveolar density between controls and mutants (Figure 3g-h). We speculate that there would be a more dramatic increase in caveolae-mediated transcytosis at a later time point due to the time required following loss of MFSD2A to change the lipid composition of brain ECs to a state permitting increased caveolae formation^8^. Importantly, besides caveolae vesicles, we have also observed morphologically diverse HRP-filled vesicles in mutant mice (Figure 3c). Thus, the increase in transcytosis in this early acute phase of attenuated canonical Wnt signaling is due to the upregulation of multiple vesicular pathways.

In summary, these data show that there is a rapid increase in transcytosis immediately following attenuation of canonical Wnt signaling in adult brain ECs, leading to BBB leakage. Notably, multiple morphologically distinct tracer-filled vesicles besides caveolae were observed at this early physiological stage, suggesting that multiple transcytosis pathways are regulated by canonical Wnt signaling in brain ECs, and expanding our understanding of the complex landscape of brain EC transcytosis regulation. The morphology of the non-caveolae vesicles observed suggests some candidate pathways, including clathrin- and Rab-mediated vesicular trafficking, uncovering potential avenues to study novel transcytosis pathway regulation at the BBB. In addition, our data show that canonical Wnt signaling regulates the molecular composition of TJs, decreasing the levels of TJ proteins in mutant mice, but that these perturbations at this early stage do not affect TJ impermeability to molecules as small as 1.9 kDa. While it is likely that longer loss of Wnt signaling would result in more severe TJ defects as well as MFSD2A-regulated caveolae-mediated transcytosis, these later time points coincide with neurological pathologies, precluding a direct, unconfounded analysis of canonical Wnt signaling regulation of brain EC subcellular properties.

Overall, these findings add to a growing body of research that demonstrates the critical importance of transcytosis for the development, maintenance, regulation, and function of the BBB in health and disease^7,31-34^. Moreover, adding onto the wide range of functions of Wnt signaling in biology, this work is able to pinpoint Wnt-mediated mechanisms that directly lead to BBB leakage – rather than secondary consequences of losing Wnt – by focusing on an early acute phase of Wnt signaling loss. Our future work, enabled by the development of more sensitive EM-compatible tools for evaluating TJ permeability, will elucidate the effect of canonical Wnt signaling on TJ permeability to small molecules at our early acute time point. These and future mechanistic insights into the maintenance of the BBB in adulthood will continue to drive the advancement of therapeutic drug delivery to the CNS, a critical need in 21^st^ century healthcare.

## Methods

### Animal Models

All mice were group housed in standard vivarium conditions, with *ad libitum* access to diet and water. *Cdh5-Cre*^*ERT2*^ mice (MGI:3848982), *Ctnnb1*^*fl/fl*^ mice (JAX:004152, MGI: J:67966), were all maintained on a C57BL/6 background. For all assays, sexually mature adult mice (8-10 weeks) of both sexes were used.

### Tamoxifen Administration

Tamoxifen (Sigma Aldrich, #T5648-1G) was dissolved in peanut oil (20 mg/ml) by brief vortexing followed by heating and rotation at 65°C and mice received five consecutive days of IP injections (0.1 mg/g body weight/day) prior to being sacrificed 3 days after the fifth injection. For the survival curve, mice received 2 mg of tamoxifen per day for 5 consecutive days.

### Immunohistochemistry

Mouse tissues were fixed by immersion in ice-cold 100% methanol (MeOH) overnight at 4°C, sequentially rehydrated in 70% MeOH/PBS, 30% MeOH/PBS, and PBS (at least 4 hrs in PBS) at 4°C, sequentially cryopreserved in 15% and 30% sucrose, and frozen in Neg-50 (Richard-Allan Scientific #6502). 20 µm tissue sections were blocked with 10% goat serum/5% BSA/PBST (0.5% Triton X-100) and stained overnight at 4°C with the following primary antibodies: α-CD31 (1:100, R&D Systems AF3628), α-ERG-488 conjugate (1:200, Abcam ab196374), α-Lef1 (1:100, Cell Signaling #2230S), α-Mfsd2a (1:100, New England Peptide, clone 9590), α-Claudin-5-488 conjugate (1:200, Thermo Fisher Scientific #352588), α-ZO-1 (1:100, Invitrogen #402200), followed by corresponding Alexa Fluor-conjugated secondary antibodies (1:250, Jackson ImmunoResearch). All sections were mounted with ProLong Gold for imaging.

### TUNEL Assay

Brains were dissected and fixed in 4% PFA/PBS overnight at 4°C, cryopreserved in 15% and 30% sucrose, and frozen in Neg-50. 20 µm brain sections underwent TUNEL staining using the Click-iT™ Plus TUNEL Assay for In Situ Apoptosis Detection, Alexa Fluor™ 647 dye (ThermoFisher Scientific #C10619). As a positive control, *Ctnnb1*^*fl/fl*^ brain slices were treated with DNase I. After the TUNEL reaction, slices were co-stained with α-ICAM2 (1:200, BD Biosciences #553326) to visualize blood vessels.

### BBB Leakage Assays

Mice were briefly anesthetized with isoflurane and EZ-Link Sulfo-NHS-LC-Biotin (100 mg/ml in PBS, 0.3 mg/g body weight) was injected into the retro-orbital sinus laterally or bi-laterally (to not exceed 100 µL in each retro-orbital sinus). After 10 minutes of circulation, brains were dissected and fixed as described above. Tissue sections were stained with α-CD31 overnight to visualize blood vessels, followed by Alexa Fluor-conjugated streptavidin (1:500, ThermoFisher Scientific #S11226) or α-Mouse IgG-conjugated secondary (1:150, ThermoFisher Scientific #A10037) for 2 hours at room temperature to visualize biotin or IgG, respectively. For HRP leakage assays, adult mice were treated as described below (see “Electron microscopy”) and slides were mounted with Permount for brightfield imaging.

### Transmission Electron Microscopy

#### Vesicular transport

To capture vesicles in a state as close as possible to their physiological dynamics, we employed low-pressure gravity perfusion, which clears the blood circulation and introduces fixative at a low flow rate equivalent to the cardiac output of adult mice, better preserving a snapshot of vesicular structures. Mice were briefly anesthetized with isoflurane and HRP type II (Sigma Aldrich, #P8250-50KU, 0.5 mg/g body weight dissolved in 0.2 ml PBS) was injected bi-laterally into the retro-orbital sinus. After 30 minutes of circulation (mice were deeply anesthetized with ketamine/xylazine after 20 minutes), mice were perfused through the heart with ∼8 mL room temperature 5% glutaraldehyde/4% PFA/0.1M sodium cacodylate buffer fixative solution (Electron Microscopy Sciences #16300, #15713-S, #11653) by low-pressure gravity-mediated flow. Brains were dissected and post-fixed in 4% PFA/0.1M sodium cacodylate buffer fixative solution overnight at 4°C. Following fixation, sagittal vibratome free-floating sections of 50 μm were collected in 0.1M sodium cacodylate buffer, incubated in 3,3’-diaminobenzidine (DAB, Sigma Aldrich #D5905-50TAB) for 35 minutes, post-fixed in 1% osmium tetroxide and 1.5% potassium ferrocyanide, dehydrated, and embedded in epoxy resin. Ultrathin sections of 70 nm were then cut from the block surface and collected on copper grids. For caveolae quantification, tissues were processed as above but without tracer injections.

#### Tight junction permeability

Mice were briefly anesthetized with isoflurane and HRP type II or microperoxidase (Sigma Aldrich #M6756-100MG, 20 mg dissolved in 0.2 ml PBS) was injected bi-laterally into the retro-orbital sinus. After 10 minutes of circulation (or 30 minutes of circulation for HRP), brains were dissected and fixed by immersion in 5% glutaraldehyde/4% PFA/0.1M sodium cacodylate buffer at room temperature for 1 hour, followed by 4% PFA/0.1M sodium cacodylate buffer at 4°C overnight. Following fixation, vibratome free-floating sections of 50 μm were collected in 0.1M sodium cacodylate buffer and incubated in 3,3’-diaminobenzidine (DAB, Sigma Aldrich #D5905-50TAB, 35 minutes for HRP, 2 hours for microperoxidase) at room temperature. As a technical note, microperoxidase was difficult to dissolve and had to be incubated in PBS for 15 minutes at room temperature (no vortexing) in order to go into solution. All tissues were processed for electron microscopy as described above.

### Microscopy, Image Processing, and Data Analysis

#### Vessel density

5 animals per genotype and 12 images per animal from the cerebellar molecular layer were used for data analysis. Images were acquired using a Leica SP8 laser scanning confocal microscope (25×, 0.75 NA) and maximum intensity projections were processed using FIJI (NIH). Vessel density was quantified as the CD31^+^ percentage area using CellProfiler.

#### LEF1

5 animals per genotype and 9 images (for cortex) or 12 images (for cerebellar molecular layer) per animal were used for data analysis, with ∼70-100 endothelial nuclei captured per image. Images were acquired using a Leica SP8 laser scanning confocal microscope (25×, 0.75 NA) and maximum intensity projections were processed using FIJI (NIH). With CellProfiler, ERG was used to segment objects (endothelial nuclei) and the LEF1 mean fluorescence intensity was quantified for each individual endothelial nucleus. For each image, the mean fluorescence intensities across all the endothelial nuclei were averaged to generate a value for each image.

#### Transmission electron microscopy

For quantification of TJ permeability, 5 animals were used per genotype and an average of 23 and 25 vessels were analyzed per animal for HRP and microperoxidase, respectively. Data were quantified as percentage of TJs impermeable to the given tracer. For HRP-filled vesicle quantification, 5 animals were used per genotype and an average of 25 vessels were analyzed per animal. For caveolae vesicle quantification, 3 or 5 animals were used per genotype and an average of 28 vessels were analyzed per animal. For HRP-filled mean vesicular density, data were quantified as the number of HRP-filled vesicles (luminal, abluminal, and cytoplasmic) per µm^2^ of endothelial cytoplasm, excluding the area of the nucleus. For caveolar density, data were quantified as the number of membrane-bound luminal or abluminal caveolae vesicles per µm of endothelial membrane. For all EM data, microvascular ECs from the cerebellar molecular layer (or choroid plexus) were used.

#### BBB leakage

12 images per animal from the cerebellar molecular layer were used for data analysis. Images were acquired using a Leica SP8 laser scanning confocal microscope (25×, 0.75 NA – only biotin and IgG) and an Olympus VS 120 slide scanner (10×, 0.4 NA for biotin and IgG, 20x, 0.4 NA for HRP). For quantification of the biotin or IgG leakage index, confocal maximum intensity projection images were processed with FIJI (NIH) and the area of the tracer was divided by the area of the CD31^+^ vessels.

#### Quantification of BBB markers

For quantification of CLDN5, ZO-1, and MSFD2A, 12 images per animal from the cerebellar molecular layer were used for data analysis. Images were acquired using a Leica SP8 laser scanning confocal microscope (25×, 0.75 NA) and maximum intensity projections were processed with FIJI (NIH). The ratio of the area of each marker to the area of CD31^+^ vessels was used to compute the expression levels and values were normalized to controls to determine the relative fold change between groups.

#### Statistical analysis

All statistical analyses were performed using Prism 7 (GraphPad Software). Two group comparisons were analyzed using an unpaired *t*-test or an unpaired *t*-test with Welch’s correction. Where appropriate, a nested *t*-test or a Mann-Whitney test was used. Multiple group comparisons were analyzed using a one-way ANOVA followed by a post-hoc Tukey’s test, a nested one-way ANOVA, or a Brown-Forsythe one-way ANOVA with Dunnett’s T3 multiple comparisons test. Sample size for all experiments was determined empirically using standards generally employed by the field, and no data was excluded when performing statistical analysis. Standard error of the mean was calculated for all experiments and displayed as errors bars in graphs. Statistical details for specific experiments, including exact *n* values, statistical tests used, and definitions of significance can be found in the figure legends.

## Supporting information

suppplemental figures

## Data Availability

The data that support the findings of this study are available from the corresponding author upon reasonable request.

## Acknowledgements

The authors thank all members of the Gu laboratory – especially Trevor Krolak, Hannah Zucker, and Joseph Amick – for comments on this manuscript; special thanks to Ralf Adams for the *Cdh5-Cre*^*ERT2*^ mice. This research was supported by an Allen Distinguished Investigator Award (C.G.), R35NS116820 (C.G.), NIH DP1NS092473 Pioneer Award (C.G.), R01 HL153261 (C.G.), RF1 DA048786 (C.G.), AHA-Allen Initiative in Brain Health and Cognitive Impairment Award (C.G.), and Fidelity Biosciences Research Initiative (C.G.). The research of C.G. was supported in part by a Faculty Scholar grant from the Howard Hughes Medical Institute. C.G. is an investigator of the Howard Hughes Medical Institute. Imaging, consultation, and/or services were in part performed in the Neurobiology Imaging Facility. This facility is supported in part by the HMS/BCH Center for Neuroscience Research as part of an NINDS P30 Core Center grant (NS072030). Electron Microscopy imaging, consultation, and services were performed in the HMS Electron Microscopy Facility. Special thanks to Maria Ericsson, Anja Nordstrom, Louise Trakimas, and Peg Coughlin for their technical assistance and support.

## Author Contributions

X.d. and C.G. conceived the project and designed experiments. X.d. performed experiments and analyzed all data. X.d. and C.G. wrote the manuscript.

