## Supplementary material for "Canonical Wnt Signaling Suppresses Brain Endothelial Cell Transcytosis to Maintain Blood-Brain Barrier Integrity": suppplemental figures

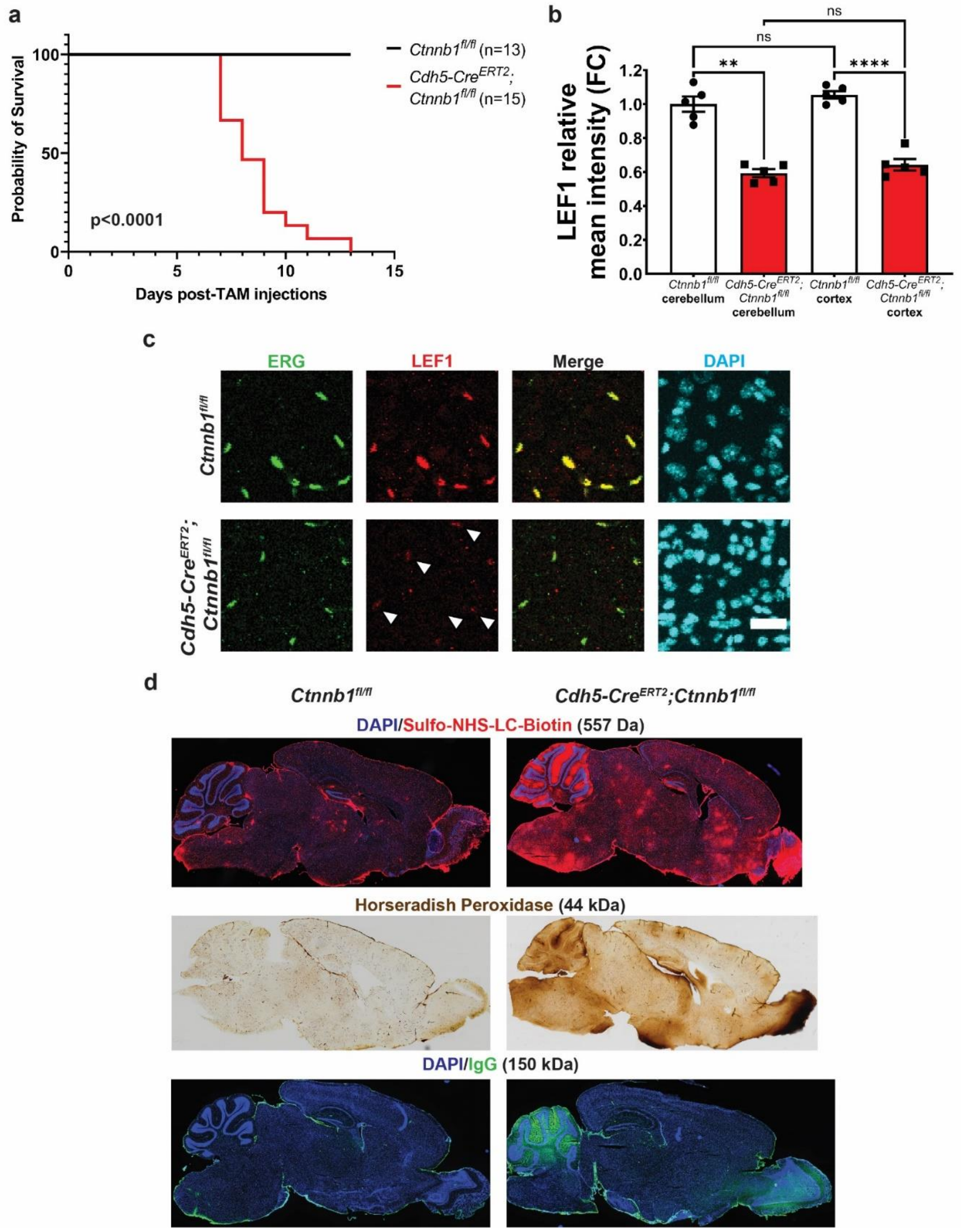


**Supplementary Figure 1. Lethality, canonical Wnt signaling attenuation, and widespread BBB leakage following acute knockout of β-catenin in adult brain ECs. (a)** Survival curve of adult control (black line) and mutant (red line) mice following five consecutive days of tamoxifen (TAM) injections at 8-10 weeks, where Day 0 represents the last TAM injection. p<0.0001 with Log-rank (Mantel Cox) test. Whereas control mice show no lethality, mutant mice begin to die 7 days after TAM injections, with 100% lethality by 13 days after injections. **(b)** Quantification of the LEF1 mean fluorescence intensity across ERG$^{+}$ EC nuclei in cerebellar molecular layer and cortical ECs (data are mean ± SEM, n=5 animals per genotype). Note that cerebellar and cortical EC nuclei in control mice have comparable levels of LEF1 and that mutant mice have the same level of LEF1 reduction in EC nuclei in the cerebellum and cortex. Cerebellar molecular layer LEF1 data are from **Figure 1f**. **p<0.01, ****p<0.0001, ns = not significant, Brown-Forsythe ANOVA with Dunnett's T3 multiple comparisons test. **(c)** Immunofluorescence images of EC nuclei (ERG), canonical Wnt signaling marker LEF1, and all nuclei (DAPI) in the cortex of control (*Ctnnb1$^{fl/fl}$*) and mutant (*Cdh5-Cre$^{ERT2}$;Ctnnb1$^{fl/fl}$*) mice. (Scale bar: 10 µm) **(d)** Immunofluorescence images of BBB leakage in control (*Ctnnb1$^{fl/fl}$*) and mutant (*Cdh5-Cre$^{ERT2}$;Ctnnb1$^{fl/fl}$*) mice: sulfo-NHS-LC-biotin (top) and IgG (bottom) with nuclear (DAPI) counterstain and brightfield image of horseradish peroxidase (middle). Low magnification images illustrate interregional heterogeneity in BBB leakage.

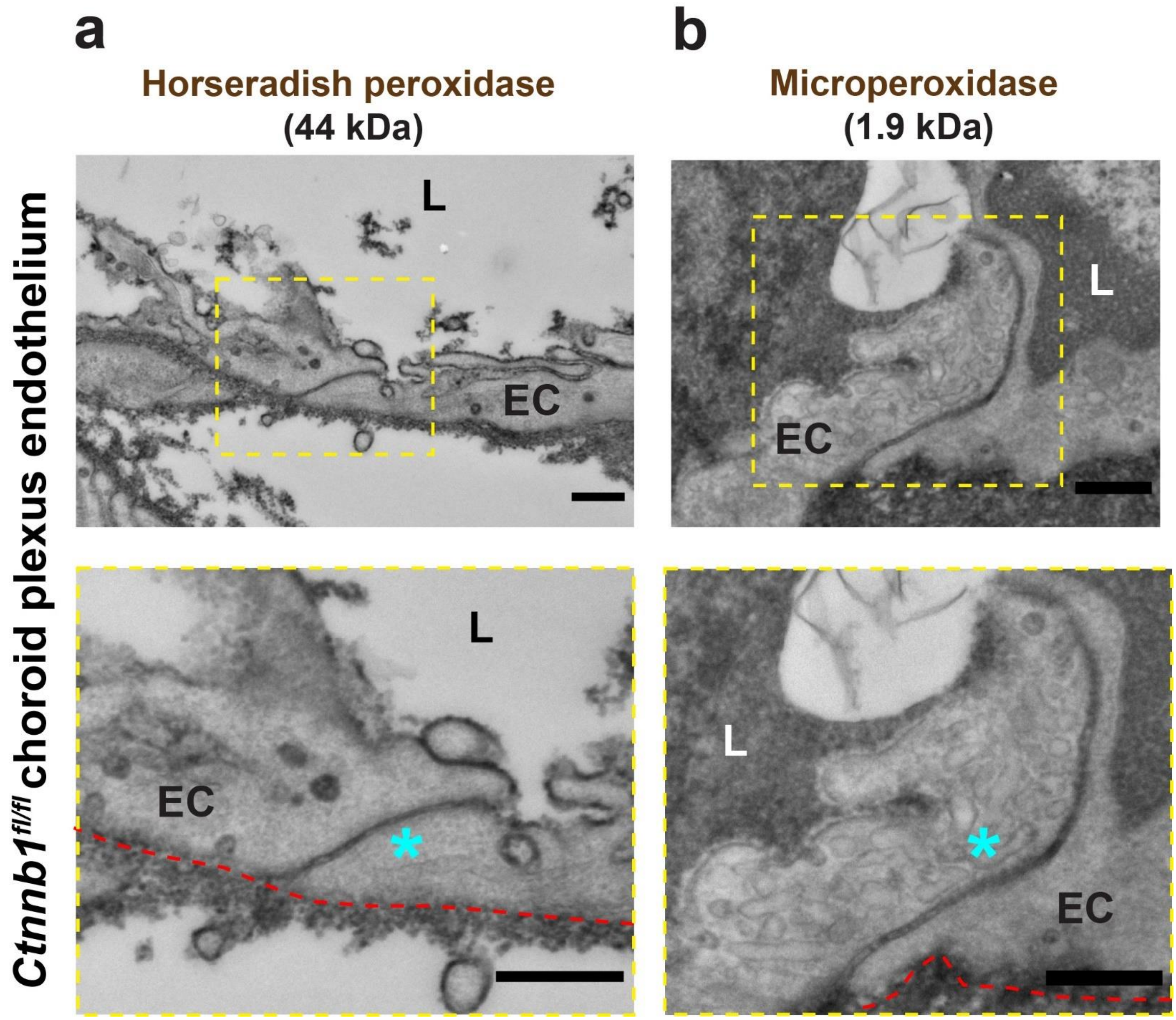


**Supplementary Figure 2. Choroid plexus microvascular endothelial tight junctions are permeable to horseradish peroxidase and microperoxidase. (a,b)** Top: Electron micrographs of choroid plexus vessel cross sections. Bottom: High magnification insets (yellow dotted line boxes from overview images) shows horseradish peroxidase (a) and microperoxidase (b) permeation through EC TJs (blue asterisk). L = lumen, EC = endothelial cell. Red dashed lines outline abluminal membranes of ECs. All scale bars: 500 nm.

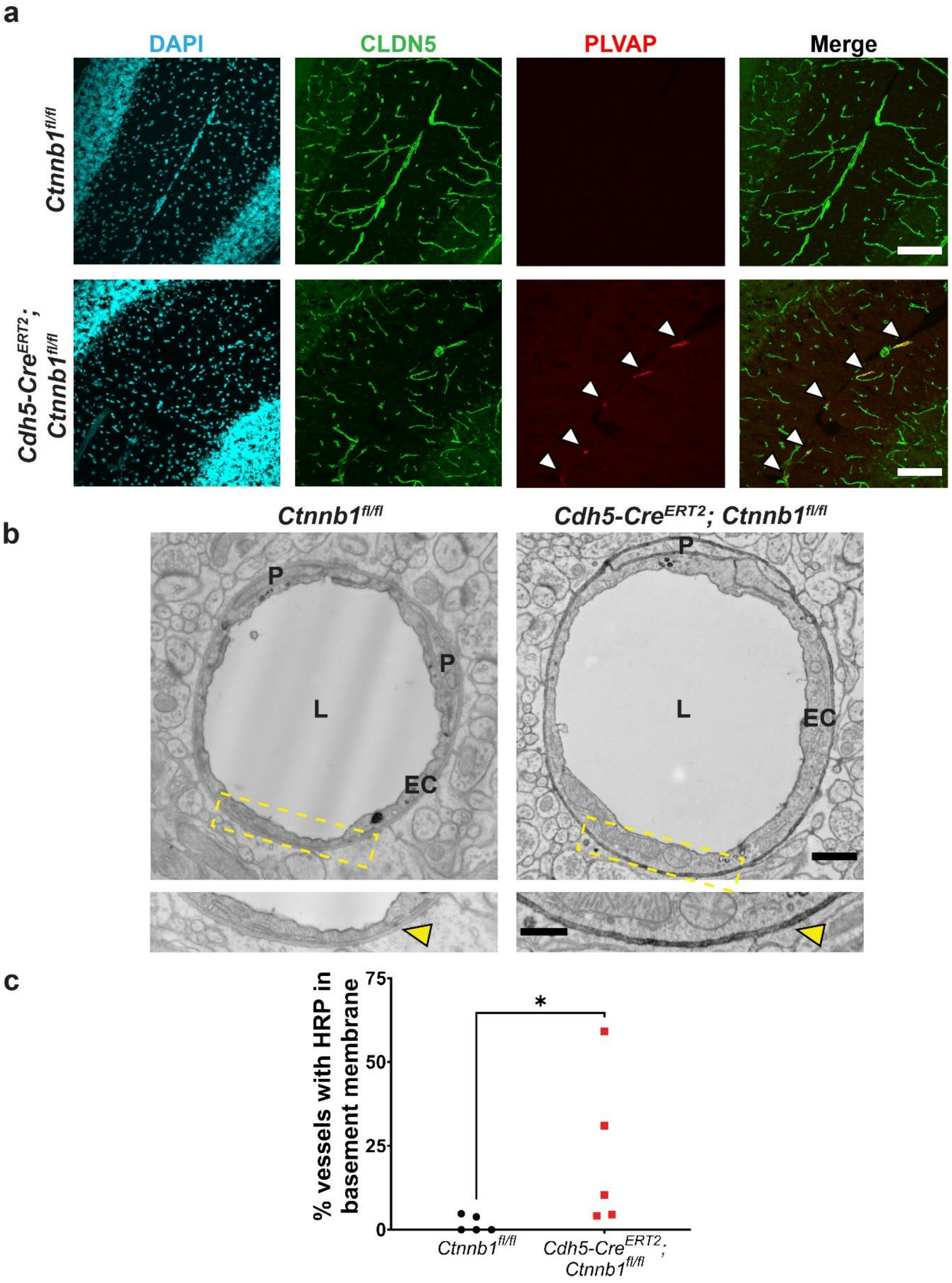


**Supplementary Figure 3. Early-phase acute attenuation of canonical Wnt signaling in the adult brain vasculature results in sparse expression of PLVAP and HRP tracer leakage into the EC basement membrane. (a)** PLVAP staining in the cerebellum with CLDN5 (vascular) and DAPI (nuclear) co-stain. White arrowheads indicate vessel segments along the central axis of the cerebellar molecular layer with sparse PLVAP expression, which also co-localize with CLDN5. Scale bar: 100 µm. **(b)** Top: Representative EM images of cerebellar molecular layer EC cross sections following HRP (black) injection and low-pressure gravity perfusion (Scale bar: 500 nm). Bottom: Insets show high magnification of areas within yellow dashed lines (Scale bar: 200 nm). Yellow arrowheads indicate HRP in the basement membrane (BM) of mutants or absence of HRP in the BM of controls. EC = endothelial cell, L = lumen, P = pericyte. **(c)** Quantification of % of vessels with HRP in BM (data are mean ± SEM, n=5 animals per genotype, *$p<0.05$, Mann-Whitney test).
